# Suberoyl bis-hydroxamic acid reactivates Kaposi’s sarcoma-associated herpesvirus through histone acetylation and induces apoptosis in lymphoma cells

**DOI:** 10.1101/2020.09.08.288837

**Authors:** Shun Iida, Sohtaro Mine, Keiji Ueda, Tadaki Suzuki, Hideki Hasegawa, Harutaka Katano

## Abstract

Kaposi’s sarcoma-associated herpesvirus (KSHV) is an etiologic agent of Kaposi’s sarcoma as well as primary effusion lymphoma (PEL), an aggressive B-cell neoplasm which mostly arises in immunocompromised individuals. At present, there is no specific treatment available for PEL and its prognosis is poor. Lytic replication of KSHV is also associated with a subset of multicentric Castleman diseases. In this study, we found that the histone deacetylase inhibitor suberoyl bis-hydroxamic acid (SBHA) induced KSHV reactivation in PEL cells in a dose-dependent manner. Next-generation sequencing analysis showed that more than 40% of all transcripts expressed in SBHA-treated PEL cells originated from the KSHV genome compared with less than 1% in untreated cells. Chromatin immunoprecipitation assays demonstrated that SBHA induced histone acetylation targeting the promoter region of the KSHV replication and transcription activator gene. However, there was no significant change in methylation status of the promoter region of this gene. In addition to its effect of KSHV reactivation, this study revealed that SBHA induces apoptosis in PEL cells in a dose-dependent manner, inducing cleavage of caspases and expression of proapoptotic factors, including Bim and Bax. These findings suggest that SBHA reactivates KSHV from latency and induces apoptosis through the mitochondrial pathway in PEL cells. Therefore, SBHA can be considered a new tool for induction of KSHV reactivation, and could provide a novel therapeutic strategy against PEL.

**Importance:** Kaposi’s sarcoma and primary effusion lymphoma cells are latently infected with Kaposi’s sarcoma-associated herpesvirus (KSHV), whereas KSHV replication is frequently observed in multicentric Castleman disease. Although KSHV replication can be induced by some chemical reagents (e.g. 12-*O*-tetradecanoylphorbol-13-acetate), the mechanism of KSHV replication is not fully understood. We found that the histone deacetylase inhibitor suberoyl bis-hydroxamic acid (SBHA) induced KSHV reactivation with high efficiency, through histone acetylation in the promoter of the replication and transcription activator gene, compared with 12-*O*-tetradecanoylphorbol-13-acetate. SBHA also induced apoptosis through the mitochondrial pathway in KSHV-infected cells, with a lower EC_50_ than measured for viral reactivation. SBHA could be used in a highly efficient replication system for KSHV *in vitro,* and as a tool to reveal the mechanism of replication and pathogenesis of KSHV. The ability of SBHA to induce apoptosis at lower levels than needed to stimulate KSHV reactivation, indicates its therapeutic potential.

## Introduction

Kaposi’s sarcoma-associated herpesvirus (KSHV, also known as human herpesvirus-8, HHV-8) is a member of gamma herpesvirus, first discovered in AIDS-associated Kaposi’s sarcoma (1). KSHV has also been identified as an etiologic agent of several lymphoproliferative disorders, including primary effusion lymphoma (PEL) and multicentric Castleman disease (2, 3). Similar to other herpesviruses, the life cycle of KSHV consists of two distinct phases; latent and lytic infection (4). Various environmental and physiological factors trigger KSHV reactivation from latency. For instance, hypoxia and reactive oxygen species have been shown to induce lytic infection (5, 6). The lytic phase is also inducible *in vitro* by some chemical reagents including 12-*O*-tetradecanoylphorbol-13-acetate (TPA) and histone deacetylase (HDAC) inhibitors (7, 8).

PEL is an aggressive B-cell lymphoma which usually presents as serous effusions without solid mass formation, arising in body cavities, such as the pleural, pericardial or peritoneal cavities (9). Most cases occur in immunocompromised individuals, such as patients with AIDS, severe immunodeficiency, or recipients of solid organ transplantation (9). Currently, no specific treatment is available for PEL and its prognosis is unfavorable (10). The median survival of PEL patients was 6.2 months under a combination of CHOP (cyclophosphamide, doxorubicin, vincristine and prednisone)-like regimen and anti-retroviral therapy, which is typically administrated for AIDS-related PEL patients (11).

HDAC inhibitors are a class of chemical compounds which exhibit various anti-tumor effects, including apoptosis, growth arrest, differentiation and autophagy (12). However, the mechanism of action is complex, as HDAC inhibitors affect gene expression profile through induction of histone acetylation (12). To date, four HDAC inhibitors have been approved by the United States Food and Drug Administration (FDA) for treatment of hematologic malignancies, although none of these are clinically available for PEL treatment (13).

In a previous study, we reported that suberoyl bis-hydroxamic acid (SBHA), a HDAC inhibitor, strongly induced KSHV lytic infection and decreased cell viability in a PEL cell line (14). However, the mechanisms underlying these effects have yet to be elucidated. In this study, we demonstrate that SBHA induced histone acetylation on the promoter region of the replication and transcription activator (RTA) gene of KSHV, resulting in KSHV reactivation. We also found that SBHA induced apoptosis of PEL cell lines through the mitochondrial pathway.

## Results

### SBHA reactivated KSHV from latency

To investigate the effect of SBHA on KSHV reactivation, two rKSHV219-infected Burkitt lymphoma cell lines (BJAB.219 and Raji.219), were exposed to various concentrations of SBHA and analyzed by flow cytometry. rKSHV219 is a recombinant KSHV which expresses red fluorescent protein (RFP) from the KSHV lytic PAN promoter and green fluorescent protein (GFP) from the EF-1a promoter (15). The analysis revealed that SBHA reactivated rKSHV.219 from latency in a dose-dependent manner (Figs. 1A and 1B). Each half maximal effective concentration (EC_50_) for rKSHV219 reactivation in BJAB.219 and Raji.219 cells was calculated as 2.95×10^-5^ M and 2.71 ×10^-5^ M, respectively (Fig. 1B). To examine whether SBHA induce KSHV reactivation in PEL cells, expression of KSHV-encoded mRNA was determined by real-time reverse transcription (RT)-PCR and protein levels was assessed by western blot. Real-time RT-PCR analysis showed that the mRNA levels of two KSHV lytic genes, RTA and viral interleukin-6 (vIL-6), were higher in SBHA-treated PEL cells than in those treated with TPA or other HDAC inhibitors (Fig. 1C). Furthermore, western blot analysis demonstrated that SBHA induced expression of RTA and vIL-6 proteins more than TPA and other HDAC inhibitors in PEL cell lines (Fig. 1D). These data indicate that SBHA strongly induces KSHV lytic infection in B-cells, including PEL cells.

**Fig. 1.**
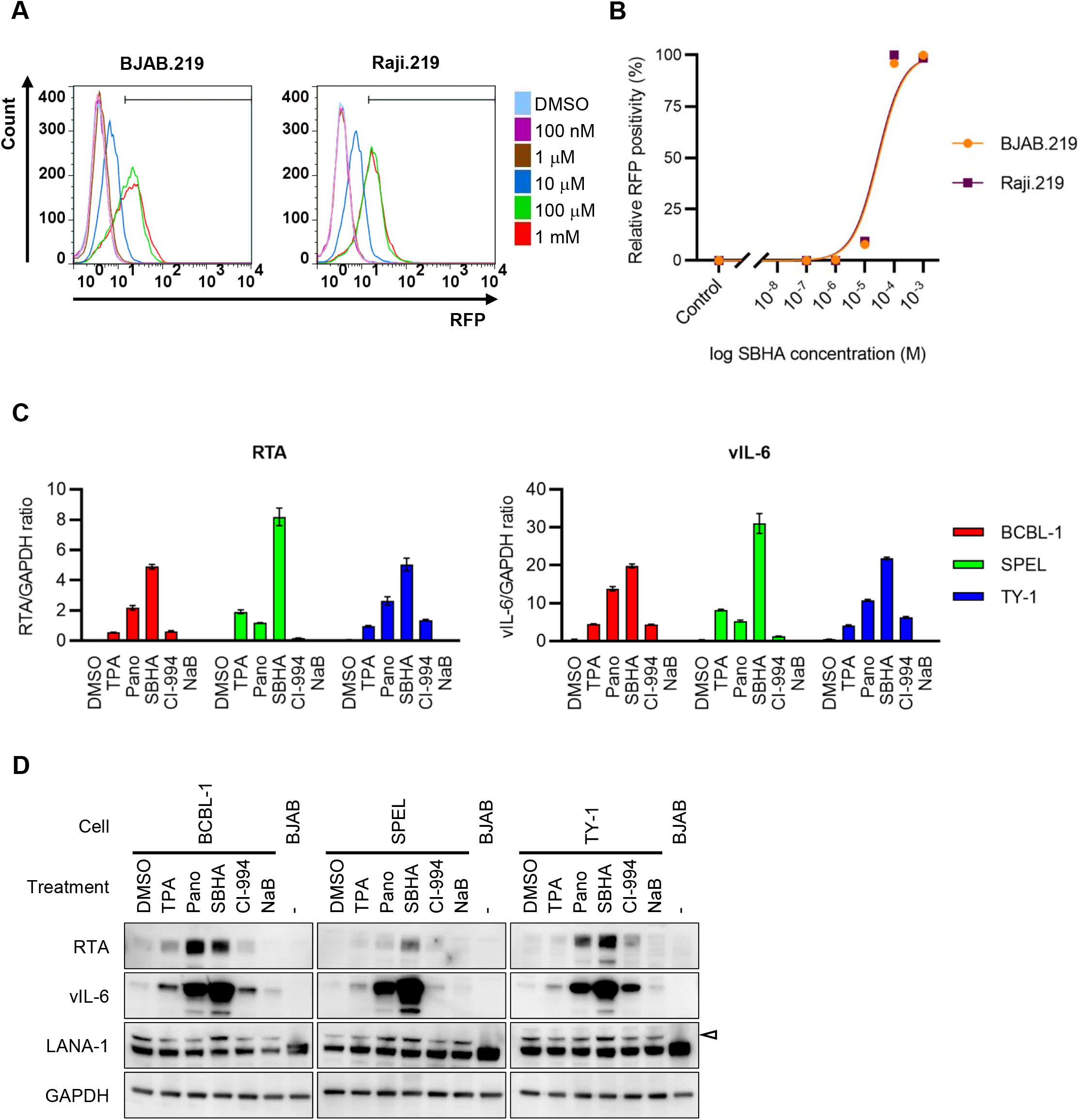
KSHV reactivation by SBHA. (A) Flow cytometry of RFP expression in rKSHV.219-infected Burkitt lymphoma cells, BJAB.219 and Raji.219. The cells were analyzed by flow cytometry 36 hours after addition of various concentration of SBHA to culture medium. (B) Dose-response relationship between SBHA concentration and RFP positivity in BJAB.219 and Raji.219 cells. (C) Real-time RT-PCR analysis for KSHV-encoded RTA (left) and vIL-6 (right) mRNA expression in three KSHV-positive PEL cell lines treated with each drug for 48 hours. The average from three independent experiments is shown and error bars indicate the standard deviation. (D) Western blot analysis for KSHV-encoded proteins, RTA, vIL-6 and LANA-1. The cells were treated with each drug for 48 hours. The arrowhead indicates LANA-1. DMSO: dimethyl sulfoxide, TPA: 12-*O*-tetradecanoylphorbol-13-acetate, Pano: panobinostat, SBHA: Suberoyl bis-hydroxamic acid, NaB: sodium butyrate.

### SBHA dramatically altered the gene expression profile of KSHV-infected PEL cells

Real-time PCR array was carried out on KSHV gene products, to evaluate the effect of SBHA on KSHV gene expression. KSHV real-time PCR array revealed the global activation of KSHV gene transcription in SBHA-treated PEL cells (Fig. 2 and supplementary Fig. 1). SBHA increased almost all KSHV gene transcripts in three different PEL cell lines, although the pattern and degree of change was very different between cell lines.

**Fig. 2.**
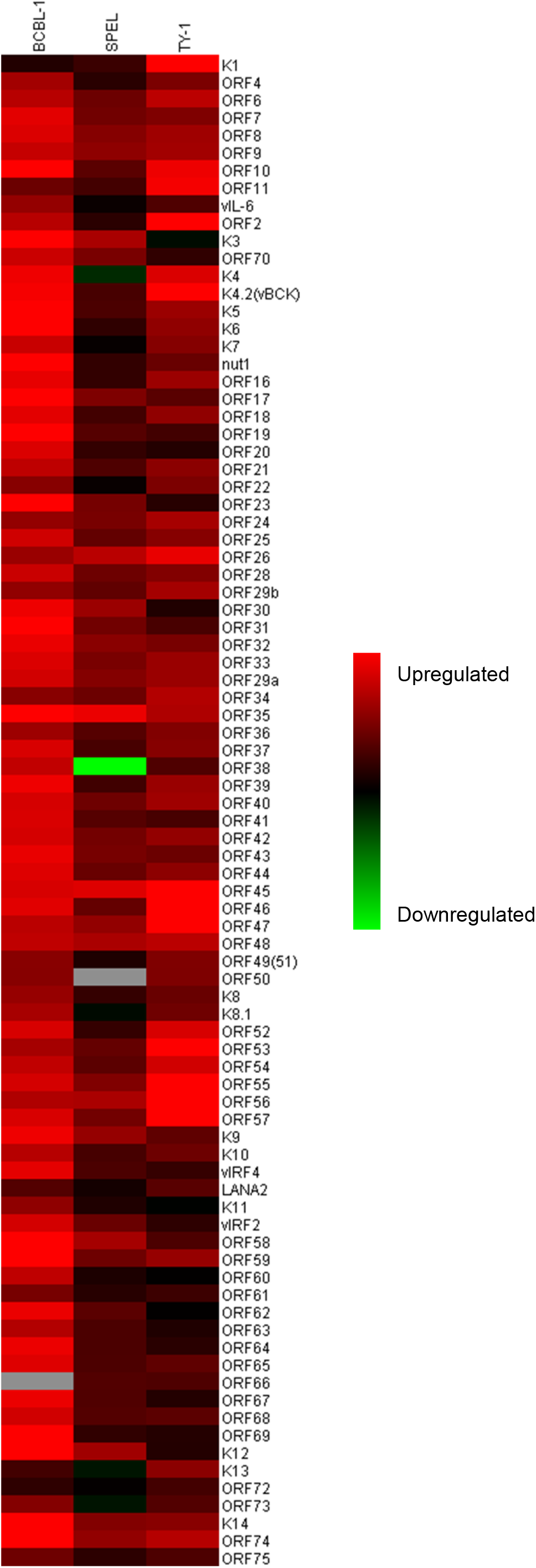
KSHV-encoded gene expression profile of SBHA treated PEL cell lines. The heatmap was generated from the results of the KSHV real-time PCR array. Red and green colors indicate upregulation and downregulation of gene transcription, respectively. Gray color indicates missing values.

Next, KSHV transcriptome analysis by next-generation sequencing was performed. As shown in Table 1, less than 1% of total reads were mapped to KSHV genome in untreated BCBL-1 and SPEL cells, and approximately 3% were mapped to KSHV genome in TPA-treated cells. In contrast, more than 40% of total reads originated from KSHV in SBHA-treated PEL cells (Table 1). Ring images of read coverage demonstrated the differences in KSHV gene expression between untreated (dimethyl sulfoxide (DMSO) only), TPA-, and SBHA-stimulated cells (Fig. 3A and supplementary Fig. 2A). The number of KSHV-mapped reads were dramatically increased by SBHA stimulation which increased expression of almost all lytic genes including vIL-6 (K2), K4, ORF45, ORF46, ORF59 and ORF65 (Fig. 3A and supplementary Fig. 2A). Notably, expression of viral interferon regulatory factors (vIRF) (including K9 (vIRF1), K10 (vIRF4), K10.5 (vIRF3) and K11 (vIRF2)) and latent genes (including K12, ORF71, ORF72 and ORF73) decreased after SBHA treatment (Fig. 3B and supplementary Fig. 2B). Taken together, these data show that SBHA intensely induces transcription of KSHV lytic genes and modifies the gene expression profile in PEL cells.

**Fig. 3.**
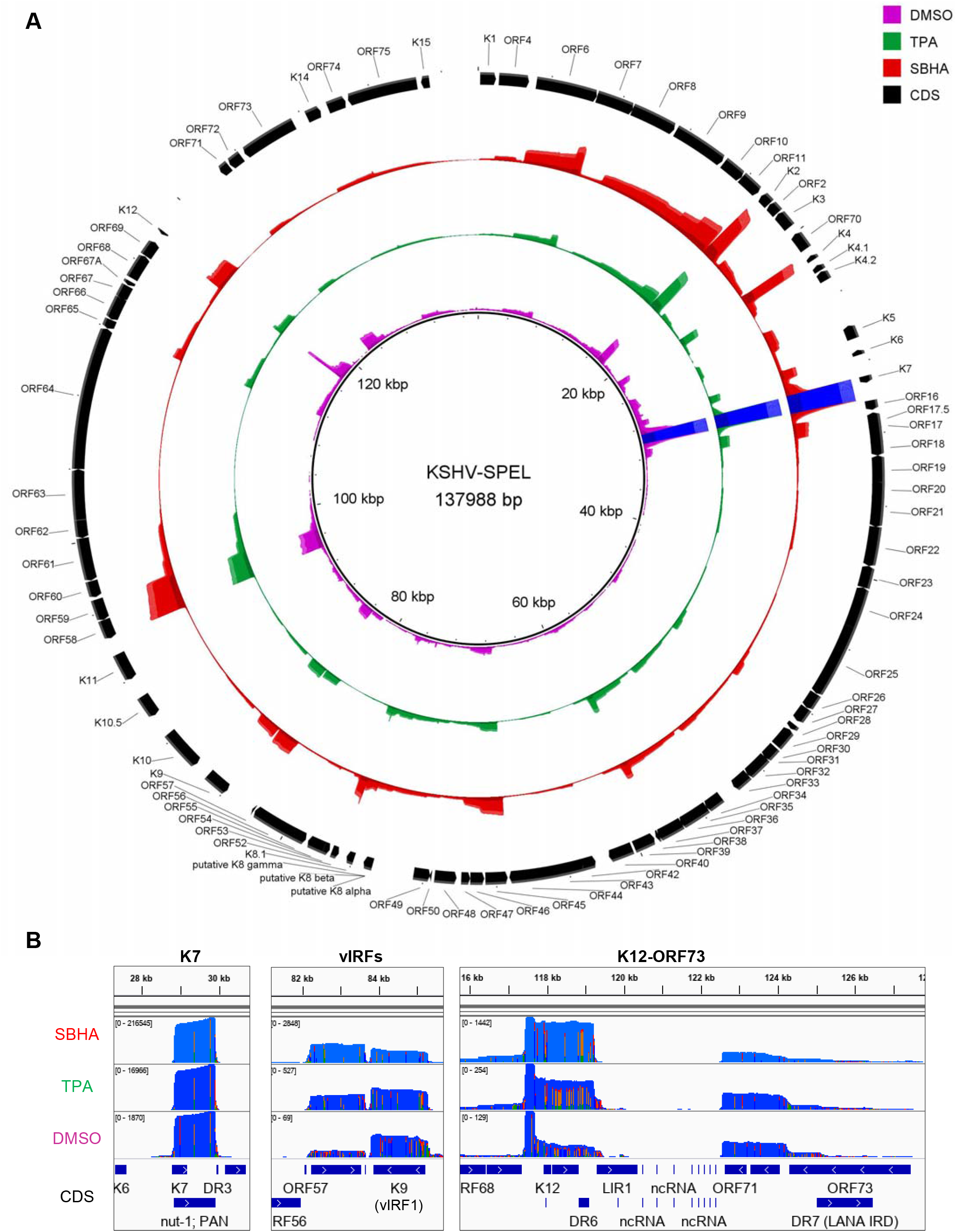
Transcriptome analysis of KSHV-encoded genes in SPEL cells. (A) Ring image for coverage of reads mapped to KSHV genome. Read coverage of DMSO (violet), TPA (green) or SBHA (red)-treated SPEL cells mapped to KSHV genome (GenBank accession no. AP017458) are shown. Maximum coverages in the image of DMSO, TPA and SBHA are 200, 1500, and 5000, respectively. Blue color in K7 indicates over the maximum reads. (B) Read coverage of SPEL cells in K7 (left), vIRF1 (center) and latent gene clusters (right).

**Table 1.**
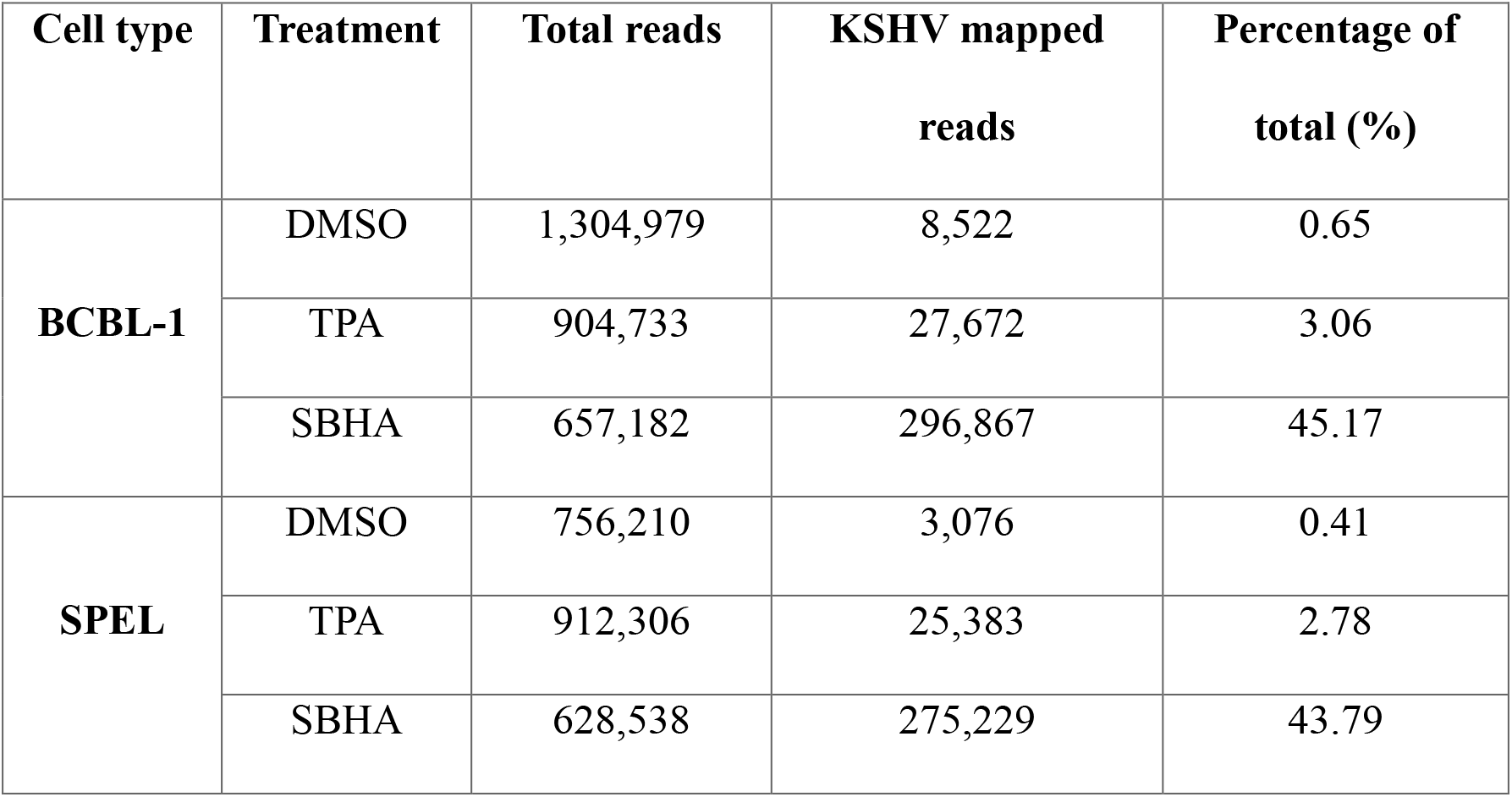
A summary of transcriptome reads mapping to the KSHV genome.

### SBHA altered histone modification but not CpG methylation of promoters of KSHV lytic genes

PEL cells were treated with TPA or SBHA for 12 hours and histone modification of the KSHV lytic gene promoter was analyzed. Chromatin immunoprecipitation (ChIP) assay revealed that activating histone modifications, acetyl-histone H3 (AcH3) and acetyl-histone H4 (AcH4), on RTA and vIL-6 promoters were strongly induced by SBHA, while minimal change was induced by TPA (Figs. 4A and 4B). The level of another activating histone mark, H3K4me3, was increased by SBHA in TY-1 and by TPA in BCBL-1. In contrast, repressive histone mark, H3K27me3, was reduced by SBHA in SPEL (Figs. 4A and 4B).

**Fig. 4.**
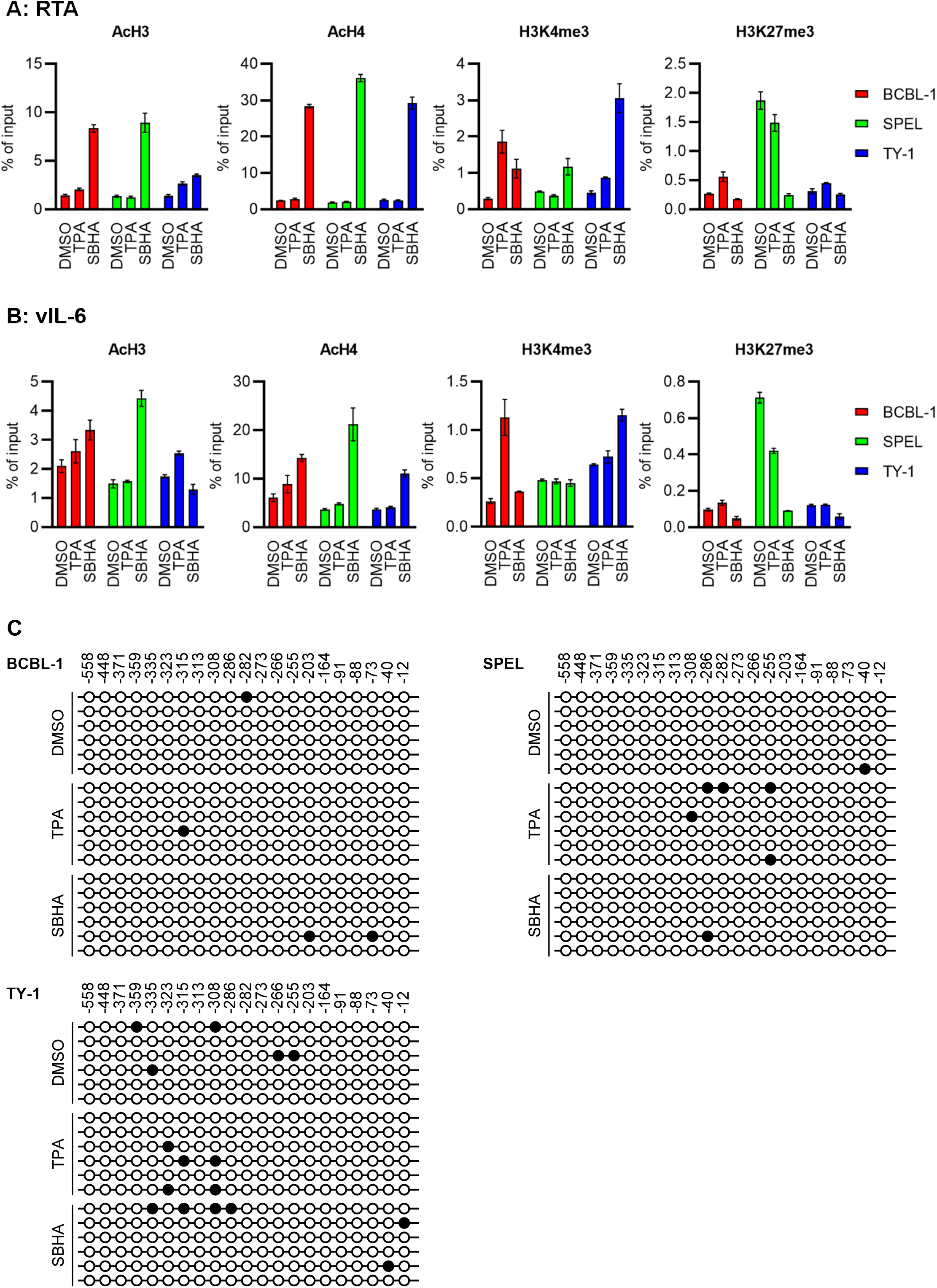
Epigenetic effects of SBHA on PEL cells. (A, B) ChIP assay for RTA (A) and vIL-6 (B) promoter. Chromatin samples were reacted with the antibody against modified histones shown at the top of each panels. The average from three independent experiments is shown and error bar indicates standard deviation. (C) Bisulfite sequencing of RTA promoter. Six clones from each cell were subjected to analysis. Closed circles indicate methylated CpG sites, while open ones indicate unmethylated CpG sites. The position relative to the transcription start site of RTA is indicated above the panel.

The methylation pattern of the RTA promoter was also investigated, since HDAC inhibitors reverse CpG methylation (16). Bisulfite sequencing analysis revealed that no or only a few CpG sites were methylated at the RTA promoter in PEL cells, regardless of SBHA or TPA treatment (Fig. 4C).

Taken together, these results indicate that reactivation of KSHV by SBHA is mainly associated with an alteration of histone modifications in the promoter of KSHV lytic genes.

### SBHA induced apoptosis and inhibited growth of PEL cells

Next, the effect of SBHA on cell growth was investigated. SBHA consistently decreased viability of PEL cell lines compared with TPA and other drugs, following 48 hours of drug treatment, demonstrated by trypan blue stains (Fig. 5A). Notably, SBHA inhibited the growth of PEL cells in a dose-dependent manner (Fig. 5B). The growth-inhibitory effects of SBHA were confirmed by XTT assay with similar results to trypan blue stain data (Figs. 5C and 5D). Each half maximal inhibitory concentration (IC_50_) for cell growth inhibition by SBHA in BCBL-1, SPEL and TY-1 cells was calculated between 2.40–5.20×10^-6^ M (Figs. 5D and 5E). The growth of Burkitt lymphoma cells (BJAB and Raji) and their recombinant KSHV-infected derivatives (BJAB.219 and Raji.219) was also inhibited by SBHA in a dose-dependent manner. However, each associated IC_50_ was higher than found in PEL cell lines (Fig 5E). Finally, Annexin V assays demonstrated that SBHA induced apoptosis in PEL cells in a dose-dependent manner (Figs. 5F and 5G). These data suggest that SBHA inhibits the growth of PEL cells by inducing apoptosis.

**Fig. 5.**
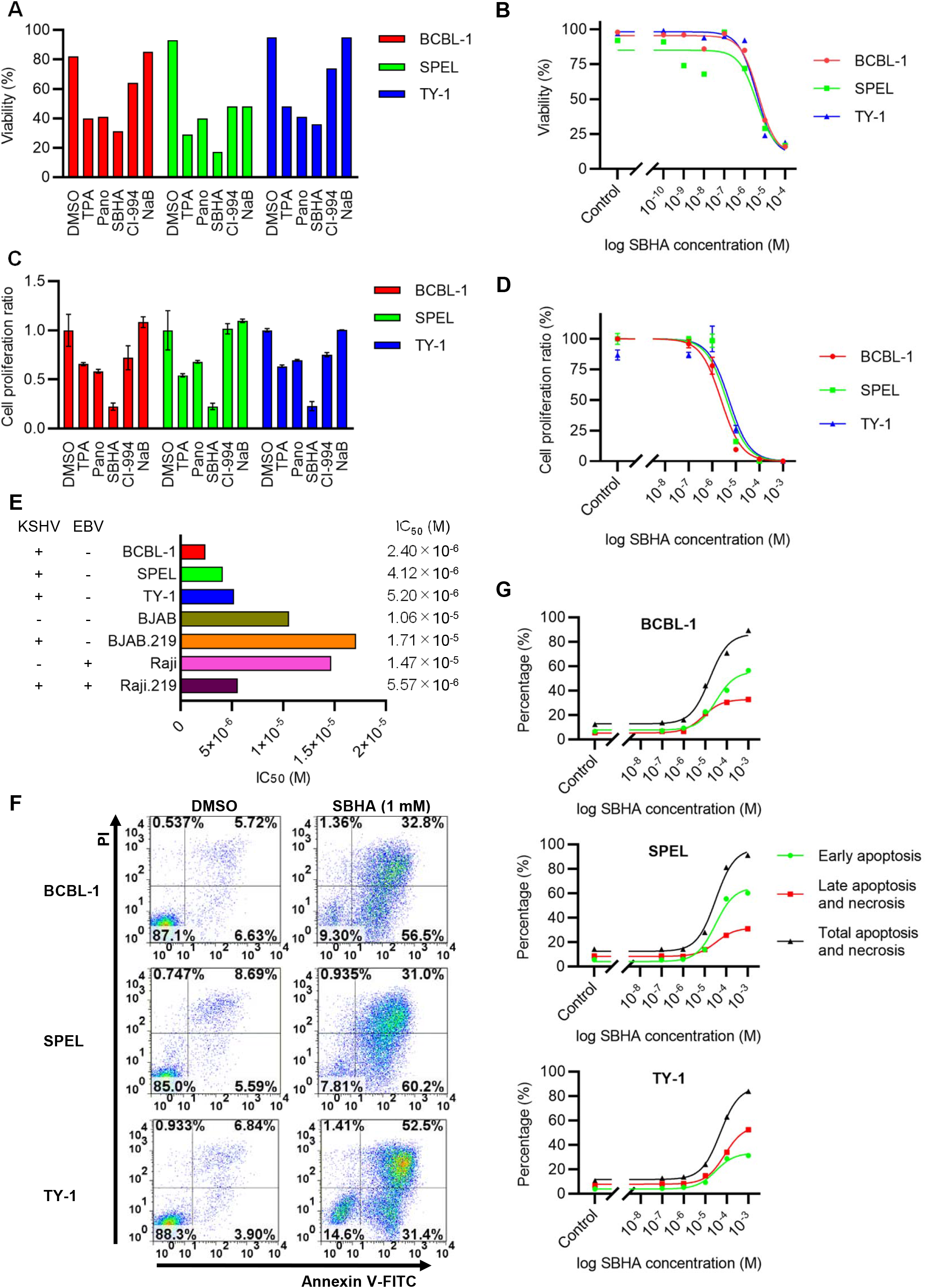
SBHA induced apoptosis in PEL cells. (A) Cell viability after drug treatment for 48 hours determined by trypan blue stain. (B) Dose-response relationship between SBHA concentration and cell viability. (C, D) XTT assay after drug treatment for 48 hours. Cell proliferation ratio was compared among TPA and several HDAC inhibitors (C), and dose-response relationship between SBHA concentration and cell growth was determined (D). The average from three independent experiments is shown and error bar indicates the standard deviation. (E) IC_50_ of each drug against various cell lines. KSHV and EBV status are shown to the left. (F) Flow cytometry of annexin V assay. Fluorescence of FITC-labeled annexin V and PI were measured following 24 hours exposure to various concentration of SBHA. The data from control (DMSO-treated) cells and SBHA-treated (1 mM) cells are shown. (G) Dose-response relationship between SBHA concentration and apoptosis of PEL cells determined by annexin V assay.

### SBHA triggered intrinsic pathway of apoptosis

The effect of SBHA on apoptosis-related factors was investigated. Western blot analysis demonstrated that expression of proapoptotic factors of the Bcl-2 family, including Bim and Bax, were upregulated by SBHA in PEL cells (Fig. 6). The cleavage of caspases downstream of Bim and Bax, including caspase-3, caspase-7 and caspase-9 was also stimulated (Fig. 6). Furthermore, SBHA downregulated Bcl-xL, an antiapoptotic factor from the Bcl-2 family. These data suggest that SBHA induces apoptosis through the mitochondrial pathway in PEL cells.

**Fig. 6.**
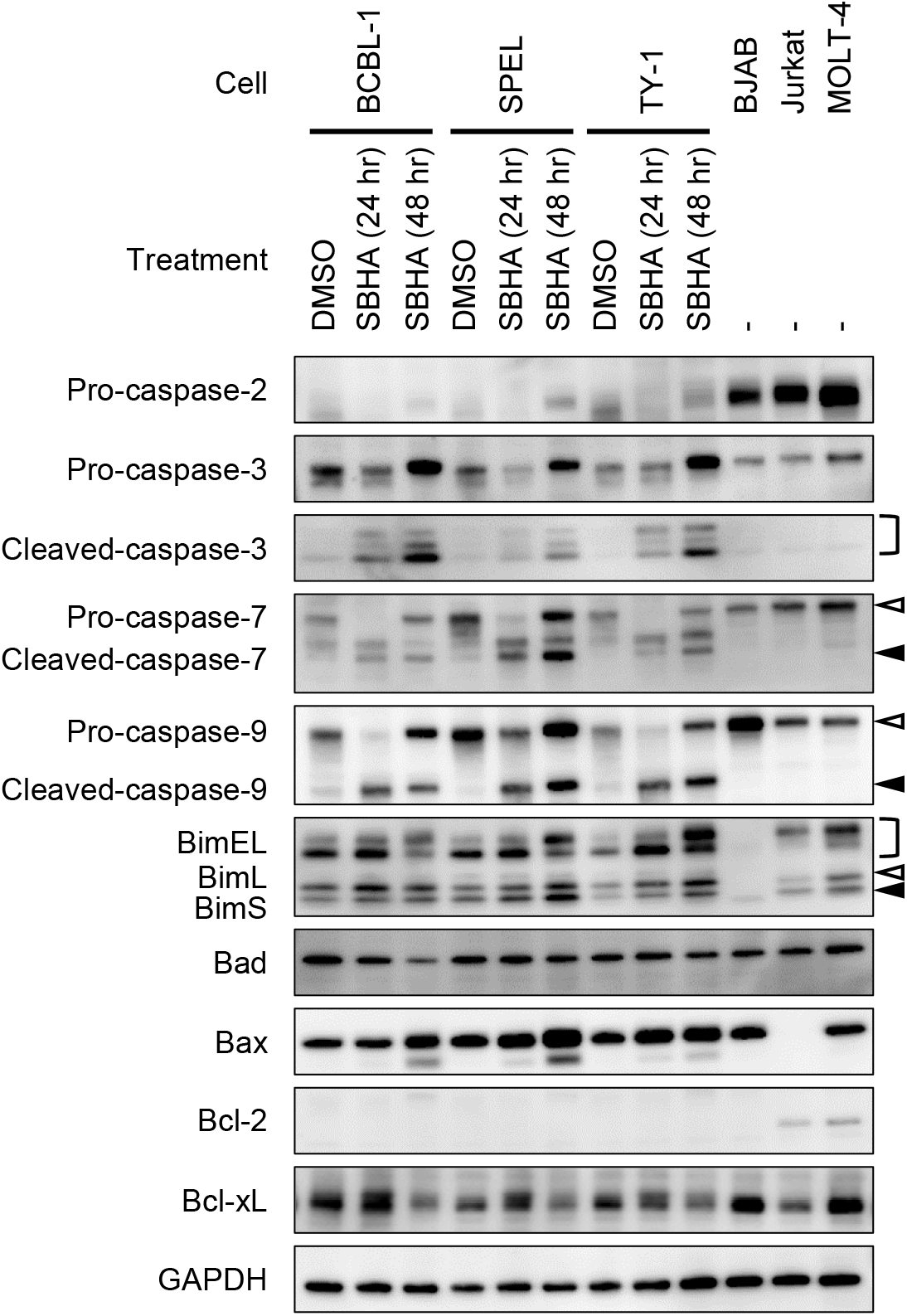
Activation of the mitochondrial pathway of apoptosis, induced by SBHA in PEL cells. PEL cells were treated with SBHA for 24 or 48 hours. Protein samples from BJAB, Jurkat and MOLT-4 cells were applied as positive control with GAPDH as loading control. White arrowheads indicate uncleaved caspases or BimL, and black arrowheads indicate cleaved caspases or BimS.

### SBHA did not stimulate Notch1 signaling pathway in PEL cells

SBHA stimulates cleavage of the full-length Notch1 protein, leading to release of the active form of Notch1, Notch1 intracellular domain (NICD), in thyroid carcinoma and neuroendocrine tumors (17–21). However, such an activation of the Notch1 signaling pathway by SBHA in PEL cells was not detected by western blot analysis (Fig. 7).

**Fig. 7.**
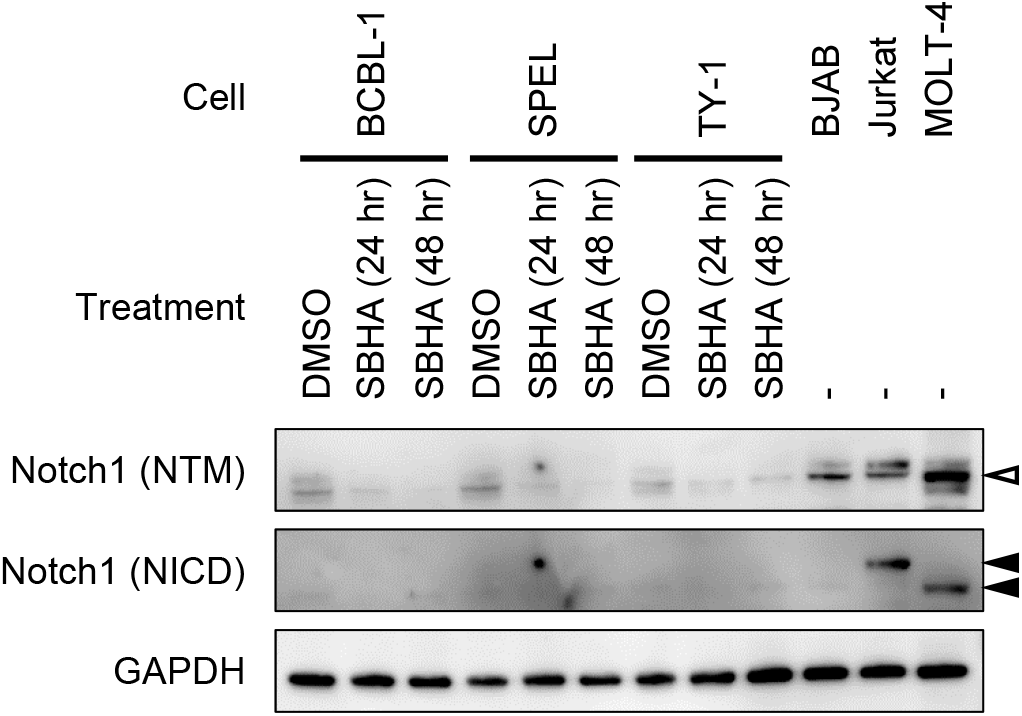
Notch1 signaling was not activated by SBHA in PEL cells. Western blot analysis for the transmembrane/intracellular region of Notch1 (NTM) and its active form, Notch1 intracellular domain (NICD), is shown. PEL cells were treated with SBHA for 24 or 48 hours. Protein samples from BJAB, Jurkat and MOLT-4 cells were applied as positive control with GAPDH as loading control. The white arrowhead indicates NTM and black arrowheads indicate NICD.

## Discussion

HDAC inhibitors are a group of chemical compounds which exhibit anti-tumor effects. Since suberoylanilide hydroxamic acid (SAHA, also known as vorinostat) was first approved for clinical use, four HDAC inhibitors have been approved by FDA for hematologic malignancies (13). SBHA, an analogue of SAHA, has been proven to demonstrate anti-tumor effects *in vitro* in various types of tumors including lung, breast and thyroid carcinomas, melanoma, acute leukemia, and myeloma (17–25). We previously showed that SBHA robustly decreased viability of SPEL cells compared with other HDAC inhibitors. To the best of our knowledge, this was the first report showing an anti-tumor effect of SBHA against a virus-associated malignancy (14). Here we demonstrate that SBHA induces apoptosis in PEL cells through the mitochondrial pathway of apoptosis and reactivates KSHV from latency by altering histone modification in the promoter region of KSHV lytic genes.

The mechanism underlying anti-tumor effect by SBHA is not yet fully understood. However, two signaling pathways have been linked to SBHA; Notch signaling and the mitochondrial pathway of apoptosis. Notch signaling is an evolutionally conserved signaling pathway which plays a crucial role in diverse developmental and physiological processes as well as tumorigenesis (26). Notch signaling is also important for pathogenesis of KSHV-associated malignancies as it induces proliferation of PEL cells and reactivation of KSHV (27, 28). Several reports have demonstrated that SBHA exhibited anti-tumor activity in neuroendocrine tumors, such as carcinoid tumor, pheochromocytoma and thyroid carcinoma, through upregulation of Notch1 signaling (17–21). However, in this study, SBHA did not activate Notch1 in PEL cells, even though trace levels of inactive Notch1 protein were detectable by western blot (Fig. 7). This variation in response to SBHA may be due to the difference in cell type or KSHV infection. In contrast to Notch signaling, the mitochondrial pathway of apoptosis is activated by SHBA in PEL cells (Fig. 6). This effect is likely based on transcriptional regulation mediated by SBHA, since expression of the proapoptotic factors Bim and Bad was upregulated. In addition, the anti-apoptotic factor Bcl-xL was downregulated by SHBA and this result accords with the hypothesis that SBHA might downregulate antiapoptotic factors through upregulating miRNAs (29). Moreover, similar upregulation of Bim and downregulation of Bcl-xL has been observed in melanoma, suggesting that activation of the mitochondrial pathway of apoptosis is the major mechanism underlying anti-tumor effect by SBHA (22).

SBHA induced histone acetylation on promoter of KSHV lytic genes, although no significant change was observed in CpG methylation of RTA promoter (Fig. 4). At present, two different groups have reported on the methylation pattern of RTA promoter, with opposite results. Chen et al. reported that CpG sites in the RTA promoter are highly methylated in BCBL-1 cells during latent infection, and that demethylation is induced by TPA treatment (30). However, Günther et al. conclude that the RTA promoter is highly methylated in HBL6 cells, but very little methylation is observed in BCBL-1 and AP3 cells in the latent phase (31). These contrasting results suggest that CpG methylation of the RTA promoter is not a key regulator for maintaining KSHV latency.

In general, lytic replication of herpesvirus leads host cell death (32). Therefore, a combination of antiviral drugs with certain drugs or stimuli which reactivate herpesvirus is regarded as a potent therapeutic tool for herpesvirus-associated malignancies; a strategy called lytic induction therapy (33). In the present study, we demonstrated that SBHA not only reactivates KSHV, but also initiates the mitochondrial pathway of apoptosis (Fig. 1 and Fig. 6). Importantly, each IC_50_ for cell growth inhibition in PEL cells (2.40×10^-6^ −5.20×10^-6^ M, Fig. 5E) is lower than the EC_50_ for KSHV reactivation (2.71×10^-5^–2.95×10^-5^ M, Fig. 1B). This suggests that the cytotoxicity of SBHA to PEL cells principally depends on the direct activation of the mitochondrial pathway of apoptosis, rather than reactivation of KSHV (Fig. 8).

**Fig. 8.**
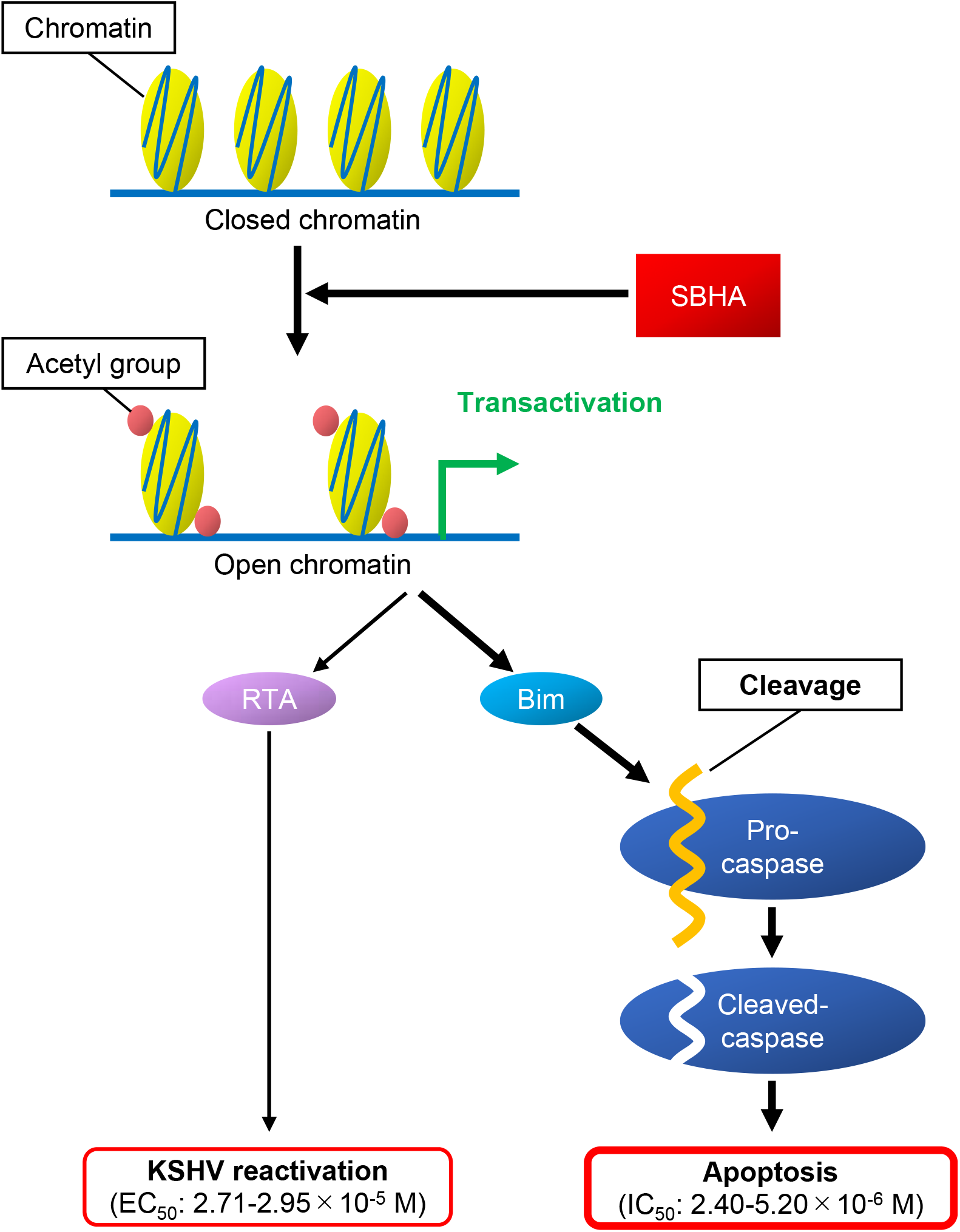
Schematic diagram for effect of SBHA in PEL cells. SBHA induces histone acetylation on promoter of various genes such as KSHV-encoded RTA and proapoptotic factors. Transactivation of target genes results in KSHV reactivation and apoptosis.

The effect of SBHA on gene expression in KSHV-infected cells was striking in that more than 40% of total transcripts expressed in SBHA-treated PEL cells were KSHV origin (Table 1). It is noteworthy that expression of vIRF genes was decreased after SBHA treatment (Fig. 3B and supplementary Fig. 2B). vIRFs interact with cellular proteins and interfere with gene transcription and signaling pathways associated with various cellular functions, such as cell death, cell cycle regulation, proliferation, and the immune response (34). According to a previous report by Muñoz-Fontela et al., vIRF3 inhibited activation of the Bim promoter by the FOXO3a transcription factor (35). Therefore, downregulation of vIRF3 transcription by SBHA may contribute to induction of Bim expression (Fig. 6).

In this study, there is little investigation into the effect of SBHA on host gene expression, including miRNAs that could contribute to the establishment of KSHV infection or tumorigenesis of KSHV-associated malignancies. However, the findings presented here provide new insight into the pathophysiology of PEL. In light of these results, SBHA should be considered a novel therapeutic strategy against PEL.

## Materials and Methods

### Cell culture and drug treatment

KSHV-positive PEL cell lines (BCBL-1, SPEL and TY-1), Burkitt lymphoma cell lines (BJAB and Raji) and T-cell leukemia cell lines (Jurkat and MOLT-4) were grown in RPMI 1640 medium supplemented with 10% fetal bovine serum at 37°C with 5% CO_2_. KSHV-infected Burkitt lymphoma cell lines (BJAB.219 and Raji.219) were established as reported previously (36): BJAB and Raji cells were infected with rKSHV.219 (kindly provided by Dr. Jeffrey Vieira, Washington University) at multiplicities of infection of 0.1 and selected in RPMI 1640 medium containing 5 μg/mL puromycin. Selected cells were maintained under the condition with 0.5 μg/mL puromycin. BJAB.219 and Raji.219 were positive for GFP expression, and KSHV genome was detected in both cells by PCR after more than 10 passages. Raji.219 is also positive for Epstein-Barr virus (EBV).

Cells were treated with TPA (Sigma-Aldrich, St Louis, MO) or one of the HDAC inhibitors; panobinostat (Sigma-Aldrich), SBHA (Sigma-Aldrich), CI-994 (Sigma-Aldrich) and sodium butyrate (Sigma-Aldrich). All reagents above were dissolved in DMSO as ×1,000 stock solution. The final concentration used for each drug was the following unless otherwise noted: 20 ng/mL for TPA, 3.6 nM for Pano, 10 μM for SBHA, 100 nM for CI-994 and 1.25 μM for NaB. Viability of cells were measured by TC10 automated cell counter (Bio-Rad, Hercules, CA) with trypan blue stain.

### Flow cytometry

To detect reactivation of KSHV by flow cytometry, red fluorescence was measured in rKSHV219-infected cells, as reported previously (15). rKSHV219-infected Burkitt lymphoma cell lines, BJAB.219 and Raji.219, were seeded at densities of 2.5×10^5^ cells/mL and incubated with SBHA at various concentrations for 36 hours. Cells were washed twice with phosphate-buffered saline (PBS) and then analyzed by CyFlow SL flow cytometer (Partec, Görlitz, Germany). Data were analyzed using FlowJo software (FlowJo, Ashland, OR).

### RNA isolation and real-time RT-PCR

Total RNA was extracted from cultured cells using RNeasy Mini Kit (Qiagen, Hilden, Germany) following the manufacture’s protocol and subsequently treated with TURBO DNase (Thermo Fisher Scientific, Waltham, MA). RNA samples were subjected to TaqMan-based real-time RT-PCR analysis to detect the mRNA of RTA, vIL-6 and glyceraldehyde-3-phosphate dehydrogenase (GAPDH) using QuantiTect Multiplex RT-PCR Kits (Qiagen) as described previously (37, 38).

### Western blot analysis

Protein extraction and immunoblotting were performed as described previously, with some modifications (39). Briefly, 1×10^6^ cells were lysed in 50 μL of M-PER Mammalian Protein Extraction Reagent (Thermo Fisher Scientific). Subsequently, 7.5 μg per lane of total protein samples were separated on 4-12% Bolt Bis-Tris Plus Gels and blotted onto polyvinylidene difluoride membrane (Bio-Rad) using the Trans-Blot Turbo Transfer System (Bio-Rad). The membranes were blocked with Block Ace solution (KAC, Kyoto, Japan) and probed with primary antibodies followed by horseradish peroxidase-conjugated anti-mouse or anti-rabbit secondary antibodies diluted with immunoreaction enhancer solution (Can Get Signal, Toyobo, Osaka, Japan). Blots were visualized by SuperSignal West Dura Extended Duration Substrate (Thermo Fisher Scientific) and signals were detected with LAS-3000 imaging system (Fujifilm, Tokyo, Japan).

The following antibodies were used in western blotting: GAPDH (sc-25778) was obtained from Santa Cruz Biotechnology (Dallas, TX). Bim (ab32158) was from Abcam (Cambridge, UK). Cleaved caspase-3 (Asp175, #9661), caspase-7 (#9492), caspase-9 (#9502), Bcl-xL (2764), Notch1 (#3608) and cleaved Notch1 (Val1744, #4147) were from Cell Signaling Technology (Danvers, MA). Apoptosis I Sampler Kit (for Bad, Bax and Bcl-2) and Apoptosis Sampler Kit II (for caspase-2 and caspase-3) were from BD Transduction Laboratories (Franklin Lakes, NJ). The mouse monoclonal antibody against RTA (40) and rabbit polyclonal antibodies against latency-associated nuclear antigen 1 (LANA-1) (41) and vIL-6 (42) were generated as previously described.

### KSHV real-time PCR array

To determine expression profiles of KSHV gene transcripts, we used KSHV real-time PCR array, a TaqMan-based real-time RT-PCR system which can detect all 87 KSHV gene transcripts simultaneously (39). RNA samples were obtained from BCBL-1, SPEL and TY-1 cells cultured with or without SBHA for 48 hours, following the procedure of RNA preparation described above. Subsequently, RNA samples were subjected to KSHV real-time PCR array using the QuantiTect Probe RT-PCR Kit (Qiagen). The copy number of each transcripts was normalized to that of GAPDH transcripts and the ratio of treated versus untreated cells was calculated.

### KSHV transcriptome analysis

BCBL-1 and SPEL cells were seeded at density of 1×10^5^ cells/mL and were stimulated with TPA or SBHA for 36 hours. Total RNA was extracted from the cells using ISOGEN (Nippon Gene, Tokyo, Japan) and then mRNA was purified from total RNA by Oligotex −dT30 Super mRNA Purification Kit (Takara Bio, Kusatsu, Japan). Double stranded cDNA was prepared from 1 μg of each mRNA sample using the PrimeScript Double Strand cDNA Synthesis Kit (Takara) and a cDNA library was established with Ligation Sequencing Kit (SQK-LSK109, Oxford Nanopore Technologies, Oxford, UK) according to the manufacturer’s instructions. Barcoded samples were pooled and ligated to sequencing adaptor. Sequencing was performed with MinION device using R9.4.1 flow cell (Oxford Nanopore Technologies).

### Bioinformatics

Guppy Version 3.6.0 was used for basecalling. Fastq files were mapped to KSHV genome (GenBank accession no. AP017458 for SPEL, and NC_003409 for BCBL-1) by CLC Genome Workbench (version 14, Qiagen), and the coverage data were obtained using Integrative Genomics Viewer (version 2.50, http://software.broadinstitute.org/software/igv/home). Ring image of the coverage was established with BLAST Ring Image Generator (version 0.95, http://brig.sourceforge.net/).

### ChIP assay

BCBL-1, SPEL and TY-1 cells were seeded at density of 5.0×10^5^ cells/mL with TPA or SBHA. After 12 hours of incubation, cells were treated with 1% formaldehyde to crosslink protein and DNA, and then unreacted formaldehyde were quenched with 0.15 M glycine. Subsequently, cells were washed three times with PBS and lysed in SDS lysis buffer (50 mM Tris-HCl, pH 8.0, 10 mM EDTA, pH 8.0, 1% SDS). Cell lysates were sonicated using Bioruptor II (Cosmo Bio, Tokyo, Japan) to shear crosslinked DNA to approximately 200–700 base pairs in length. Chromatin samples derived from 1.0×10^6^ cells were precleared with protein G sepharose (GE Healthcare, Chicago, IL) and then 1-5 μg of each antibody was added to form immune complex. After overnight incubation, immune complexes were precipitated with protein G sepharose and sequentially washed with low salt immune complex wash buffer (0.1% SDS, 1% Triton X-100, 2 mM EDTA, pH 8.0, 20 mM Tris-HCl, pH 8.0, 150 mM NaCl), high salt immune complex wash buffer (0.1% SDS, 1% Triton X-100, 2 mM EDTA, pH 8.0, 20 mM Tris-HCl, pH 8.0, 500 mM NaCl), LiCl immune complex wash buffer (250 mM LiCl, 1% NP-40, 1% sodium deoxycholate, 1 mM EDTA, pH 8.0, 10 mM Tris-HCl, pH 8.0) and TE buffer. Finally, protein/DNA complexes were eluted with elution buffer (1% SDS, 100 mM NaHCO_3_), incubated with 0.2 M NaCl at 65°C overnight to reverse crosslink, and then treated with RNase A and proteinase K. For preparation of input DNA, chromatin samples from 1.0×10^4^ cells were sequentially treated with 0.2 M NaCl, RNase A and proteinase K. All DNA samples were subjected to real-time PCR analysis using QuantiTect SYBR Green PCR Kit (Qiagen) with primers shown below.

The following antibodies were used for chromatin immunoprecipitation: Normal rabbit IgG (PM035) was obtained from Medical & Biological Laboratories (Nagoya, Japan). Acetyl-histone H3 (#06-599), acetyl-histone H4 (#06-866), trimethyl-histone H3 (Lys4, #04-745) and trimethyl-histone H3 (Lys27, #07-449) were from Millipore (Burlington, MA). The sequences of the primers were as follows: RTA promoter, forward primer 5’-GGTACCGAATGCCACAATCTGTGCCCT-3’, reverse primer 5’-ATGGTTTGTGGCTGCCTGGACAGTATTC-3’ (43); vIL-6 promoter, forward primer 5’-GCGCCTCCCGGTACAAGTCC-3’, reverse primer 5’-GACCATTGGCGGGTAGAATC-3’ (44).

### Bisulfite sequencing

BCBL-1, SPEL and TY-1 cells were cultured with TPA or SBHA for 24 hours. DNA samples were prepared from the cells using DNeasy Blood & Tissue Kit (Qiagen). Before bisulfite conversion, 5 μg of each DNA sample was digested with EcoRI. Digested DNA was denatured in 0.3 M NaOH and subjected to bisulfite conversion with 4.0 M sodium bisulfite/0.4 mM hydroquinone (pH 5.0) at 55°C for three hours. Bisulfite-converted DNA was sequentially treated with 0.3 M NaOH and 0.45 M ammonium acetate for deamination and desulfonation, respectively. Purified DNA was subjected to PCR to amplify the promoter region of the RTA gene using KOD −Multi & Epi-enzyme (Toyobo, Osaka, Japan) with the following primers: forward primer 5’-GTGTTTTATTATTTTTATAG-3’, reverse primer 5’-CATCTAACATAACTTTAATC-3’ (31). PCR products were cloned into the pCR2.1 vector using the TA Cloning Kit (Invitrogen, Carlsbad, CA) to isolate single clones. The DNA sequence of each clone was determined by Sanger sequencing using M13 forward and reverse primers.

### XTT assay

XTT assays were performed using Cell Proliferation Kit II (XTT) (Roche Diagnostics, Mannheim, Germany) following manufacturer’s protocol. Briefly, cells were seeded into 96-well plates at a density of 1×10^5^ cells/mL with TPA or a HDAC inhibitor. After 48 hours of culture, XTT labeling mixture was added to each well. Following a further four hours incubation, absorbance at 450 nm was measured by iMARK microplate reader (Bio-Rad).

### Annexin V assay

Annexin V assay was carried out using MEBCYTO Apoptosis Kit (Annexin V-FITC Kit) (Medical & Biological Laboratories) according to manufacturer’s instructions. In brief, cells were seeded at density of 5.0×10^5^ cells/mL and incubated with various concentration of SBHA for 24 hours to induce apoptosis. After washing with PBS, the cells were stained by fluorescein isothiocyanate (FITC) labeled annexin V and propidium iodide (PI). Finally, cells were analyzed by a flow cytometer as described above.

### Data analysis

Generation of dose-response curves, and calculation of EC_50_ and IC_50_ were carried out using GraphPad Prism 8.3.0 software (GraphPad Software, San Diego, CA). Data obtained from KSHV real-time PCR arrays were analyzed using Cluster 3.0 software (45) and heatmaps were generated using Java Treeview software (46).

### Accession numbers

The sequence reads by NGS in this study were registered in Sequence Read Archive database as accession number DRA010668 in the DNA Data Bank of Japan Sequence Read Archive.

## Acknowledgments

The authors thank Dr. Jeffrey Vieira, Department of Laboratory Medicine, University of Washington, for providing the recombinant KSHV. This work was partly supported by the Japan Society for the Promotion of Science (grant no. 19K07600 to HK), and Japan Agency for Medical Research and Development (AMED, no. JP20fk0410016 to HK and JP20fk0108104 to TS and HK).

## Supplementary data

**Sup Fig. 1.**
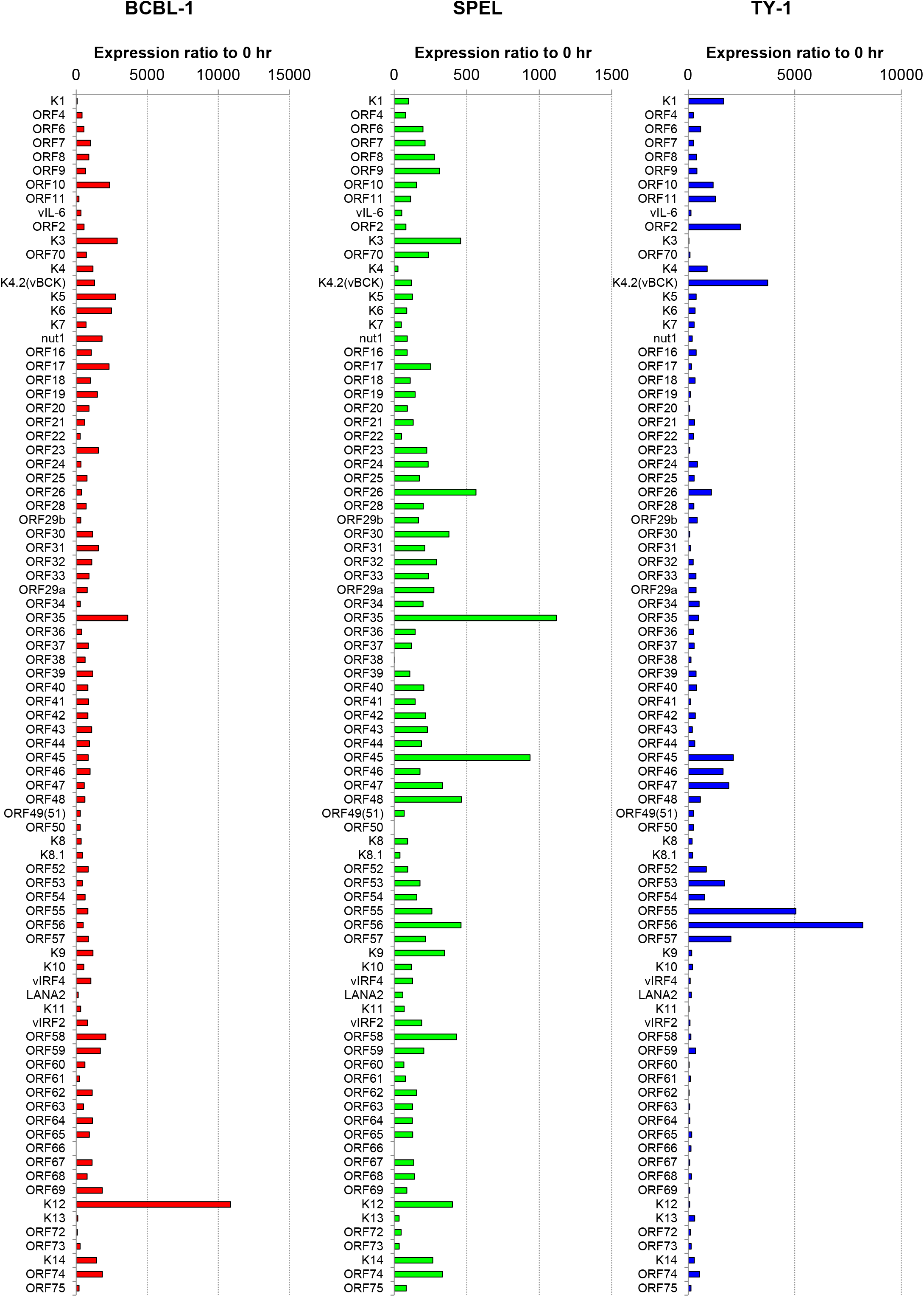
Alteration of KSHV-encoded gene expression induced by SBHA. PEL cells were treated with SBHA for 48 hours and RNA samples were subjected to KSHV real-time PCR array to determine expression profiles of KSHV-encoded genes. The copy number of each transcript was normalized to that of GAPDH and the ratio to the value of untreated cells is shown.

**Sup Fig. 2.**
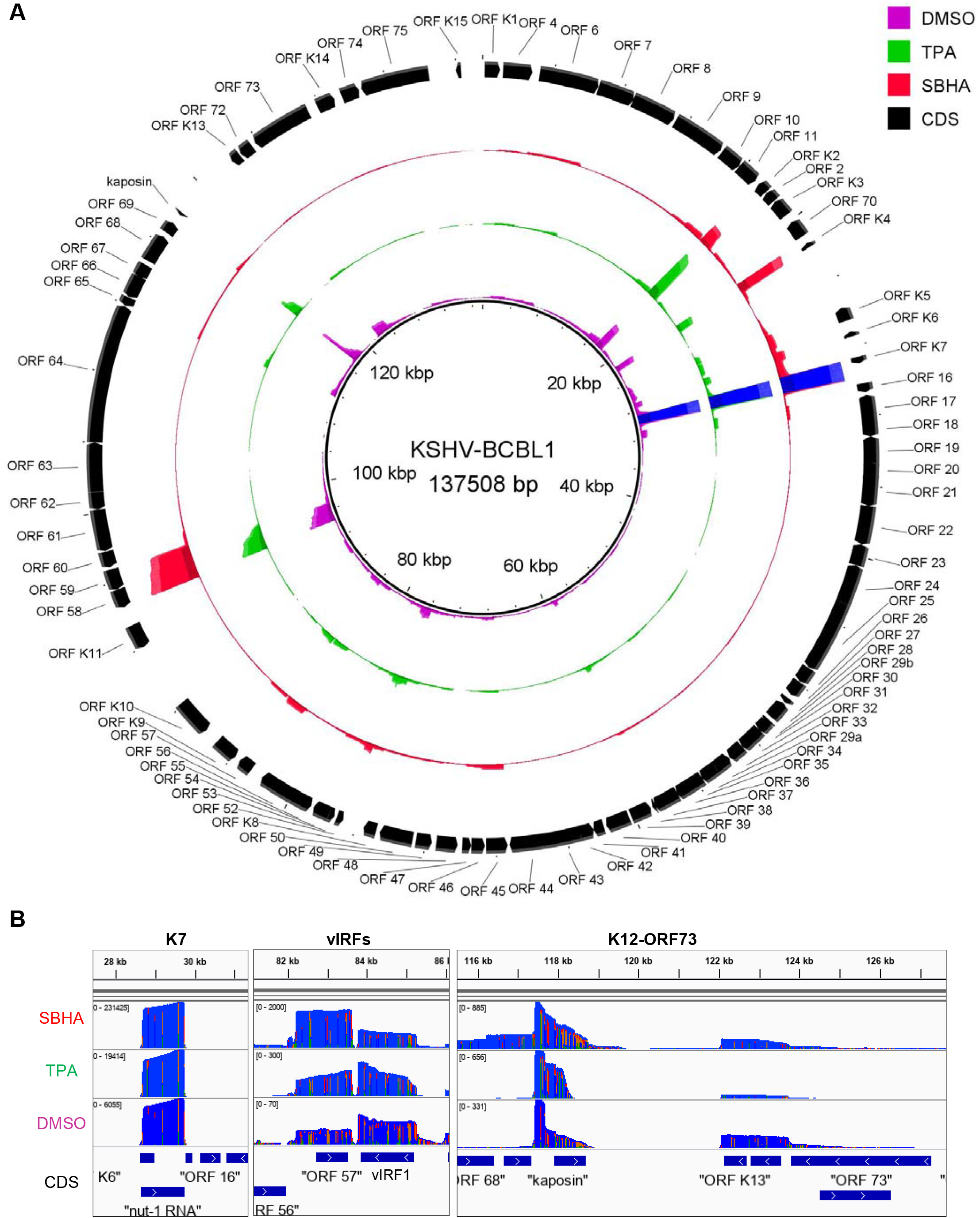
Transcriptome analysis of KSHV-encoded genes in BCBL-1 cells. (A) Ring image for coverage of reads mapped to KSHV genome. Read coverage of DMSO (violet), TPA (green) or SBHA (red)-treated BCBL-1 cells mapped to KSHV genome (GenBank accession no. NC_003409) are shown. Maximum coverages in the image of DMSO, TPA and SBHA are 500, 2000, and 15,000, respectively. Blue color in K7 indicates over the maximum reads. (B) Read coverage of BCBL-1 cells in K7 (left), vIRF (center) and latent gene clusters (right). Coverage of SBHA (top), TPA (2nd), and DMSO (3rd)-treated cells are shown. The lowest column indicates coding sequences of open reading frame.

